# Amyloid-β and tau pathologies relate to distinctive brain dysconnectomics in preclinical autosomal dominant Alzheimer’s disease

**DOI:** 10.1101/2021.07.03.450933

**Authors:** Edmarie Guzmán-Vélez, Ibai Diez, Dorothee Schoemaker, Enmanuelle Pardilla-Delgado, Clara Vila-Castelar, Joshua T. Fox-Fuller, Ana Baena, Reisa A. Sperling, Keith A. Johnson, Francisco Lopera, Jorge Sepulcre, Yakeel T. Quiroz

## Abstract

The human brain is composed of functional networks that have a modular topology, where brain regions are organized into communities that form internally dense (segregated) and externally sparse (integrated) subnetworks that underlie higher-order cognitive functioning. It is hypothesized that amyloid-β and tau pathology in preclinical Alzheimer’s disease (AD) spread through functional networks, disrupting neural communication that results in cognitive dysfunction. We used high-resolution (voxel-level) graph-based network analyses to test whether *in vivo* amyloid-β and tau burden was associated with the segregation and integration of brain functional connections, and episodic memory, in cognitively-unimpaired Presenilin-1 E280A carriers who are expected to develop early-onset AD dementia in approximately 13 years on average. Compared to non-carriers, mutation carriers exhibited less functional segregation and integration in posterior default-mode network (DMN) regions, particularly the precuneus, and in the retrospenial cortex, which has been shown to link medial temporal regions and cortical regions of the DMN. Mutation carriers also showed greater functional segregation and integration in regions connected to the salience network, including the striatum and thalamus. Greater tau burden was associated with lower segregated and integrated functional connectivity of DMN regions, particularly the precuneus and medial prefrontal cortex. In turn, greater tau pathology was related to higher segregated and integrated functional connectivity in the retrospenial cortex, and the anterior cingulate cortex, a hub of the salience network. These findings enlighten our understanding of how AD-related pathology distinctly alters the brain’s functional architecture in the preclinical stage, possibly contributing to pathology propagation and ultimately resulting in dementia.

**SIGNIFICANCE STATEMENT:** Amyloid-β and tau pathologies in Alzheimer’s disease (AD) are hypothesized to propagate through functional networks. Research is needed to elucidate how AD-related pathologies alter the brain’s functional connections years before symptom-onset to help predict pathology accumulation and dementia risk. Using high-resolution (voxel-level) graph-based network analyses and PET imaging, we showed that greater tau burden was associated with largely dysconnectivity of posterior default-mode network brain regions, and increased connectivity of structures associated with the salience network and others that are critical for integrating information across neural systems in cognitively-unimpaired Presenilin-1 E280A carriers. Findings enlighten how amyloid-β and tau relate to distinct patterns of functional connectivity in regions that underlie memory functioning and are critical for information processing, helping predict disease progression.

## INTRODUCTION

The progressive accumulation of amyloid-β plaques and neurofibrillary tangles in Alzheimer’s disease (AD) begin several years before symptom onset (i.e., the preclinical stage)(1, 2) and follows distinct spatio-temporal patterns(3, 4). Amyloid-β accumulates throughout anatomically distant neocortical regions two decades before the onset of memory impairment(2, 5), whereas tau accumulation is first observed in the transentorhinal and entorhinal cortex, later spreading into adjacent association and unimodal cortices(6–8). These patterns of pathology accumulation are highly reminiscent of functional brain networks, which can be represented by the interregional coherence of spontaneous neural activity fluctuations during rest using functional magnetic resonance imaging (fMRI). This has led some to hypothesize that pathology propagation in AD may occur through functional networks(9–11). *In vivo* neuroimaging studies have shown that amyloid-β first accumulates in a set of distributed brain regions that form part of the so-called default mode network (DMN) (i.e., precuneus, medial prefrontal and inferior parietal cortices), considered a backbone of mainly cortical integration(12, 13). Conversely, tau pathology begins to accumulate in vulnerable *loci*, later advancing to local functionally connected regions, possibly following trans-synaptic spread along functional connections(8, 9, 14). As such, characterizing how AD-related pathology relates to the brain’s functional architecture in the preclinical stage of AD can improve our understanding of the impact of pathology on neural communication and propagation early in the disease, and help with early detection, disease staging and prediction of cognitive decline.

Some functional brain networks have been shown to undergo two distinct phases that are differentially associated with AD stages. Some studies have reported increased functional connectivity (i.e., hyperconnectivity) within the DMN in cognitively-unimpaired older adults with high levels of amyloid-β but low tau, and reduced functional connectivity (i.e. hypoconnectivity) in those with high levels of both amyloid-β and tau(15–17). Some DMN regions, particularly the precuneus, have also been shown to act as mediators between hyperconnected and hypoconnected hubs(16), which are brain regions that are highly interconnected and that act as way stations to integrate information across different and often segregated or distributed neural systems. While some studies have suggested an inverse relationship between tau deposition and functional connectivity(16, 18), others have shown that elevated tau deposition is associated with stronger local functional connectivity early in AD(10, 19, 20). Greater functional connectivity in tau pathology hotspots have been related to a faster accumulation of tau in interconnected regions, suggesting that functional connectivity in regions that accumulate tau faster and earlier may enable the spread of tau to nearby structures that are closely functionally connected(10, 19, 21).

These complex brain networks have a modular organization — i.e., different voxels of the brain are organized in communities that specialize on distinct tasks. These modules have a large number of links that connect voxels within a community (network segregation) and a smaller set of links that integrate information between communities (network integration). However, very little is known about how early AD-related pathology—particularly tau pathology, a strong predictor of neurodegeneration and cognitive decline(22–24)—distinctly impact the integration and segregation of networks crucial for efficient information processing. Most of the published research has been limited by focusing on amyloid-β and not tau, and by studying older adults who are at high risk but may not develop AD-dementia. Further, many studies have examined functional connectivity by averaging sets of pair-wise correlations, thereby obscuring the intricate dynamics of functional networks and potential effects on cognitive functioning in the preclinical stage of AD.

To address these shortcomings in the field, we used high-resolution (voxel-level) graph-based network analyses to characterize the topological organization of functional connections in the brain associated with *in vivo* amyloid-β and tau burden, and episodic memory in individuals with autosomal-dominant AD (ADAD) caused by the Presenilin-1 *(PSEN1)* E280A mutation. Studying ADAD provides a unique opportunity to understand the associations between AD-related pathology and functional connections in preclinical AD, as mutation carriers are virtually guaranteed to develop dementia and have limited age-related comorbidities that could contribute to brain dysfunction. *PSEN1* E280A carriers have a well-characterized clinical profile, with an onset of mild cognitive impairment (MCI) at a median age of 44 years and dementia at 49 years (25). They exhibit widespread cortical amyloid-β accumulation approximately two decades before estimated MCI onset, and tau pathology in the entorhinal cortex close to six years before estimated MCI onset(2, 24). They also show reduced glucose metabolism in temporal and parietal regions, including in the precuneus, more than a decade before estimated MCI onset(26).

We computed a data-driven voxel-level whole-brain analysis to create maps of functional hubs that help characterize segregated (within-network) and integrated (between-network) connections and examined whether the segregation and integration of functional networks at resting differed between mutation carriers and age-matched non-carrier family members, and their relationship to AD-related pathology burden and episodic memory. Amyloid-β and tau burden were measured with 11C Pittsburg compound B and [F18] Flortaucipir Positron Emission Tomography (PET) respectively. We hypothesized that there would be significant group differences in the segregation and integration of functional connections in DMN regions, particularly in the precuneus and medial temporal regions. We also hypothesized that less segregated functional connectivity would be associated with greater tau burden and worse episodic memory, and stronger integrated functional connectivity with higher amyloid-β burden and worse episodic memory.

## RESULTS

### Sample characteristics

We examined 21 cognitively-unimpaired *PSEN1* E280A carriers who were on average 8 years from expected symptom onset and 13 years from expected dementia onset, and 31 non-carriers. Groups did not significantly differ in age, education, sex, or Mini-Mental State Examination (MMSE) scores (Table 1). Compared to non-carriers, mutation carriers performed significantly worse than non-carriers in the Consortium to Establish a Registry for Alzheimer’s Disease (CERAD) word list delayed recall, a measure of episodic memory that is sensitive to early cognitive changes in this population (*F* (1,49) =7.87, p=.007, *d*=0.73) (Table 1)(25).

**Table 1.** Demographic and cognitive data.

|  | Non-carriers<br>(n=31) | Mutation Carriers<br>(n = 21)<br>*M (SD)* | *p* |
| --- | --- | --- | --- |
| Demographics |  |  |  |
| Age (years) | 35.59 (4.75) | 35.99 (5.02) | .775 |
| Education | 11.68 (4.00) | 10.19 (3.84) | .185 |
| Sex (% males) | 48.4 | 47.7 | .957 |
| Cognitive tests |  |  |  |
| FAST (n scores ½) | 29/4 | 16/5 | .072 |
| MMSE | 28.9 (.91) | 28.43 (.98) | .085 |
| CERAD delayed recall | 7.81 (1.28) | 6.52 (2.06) | .007 |

*Note.* M = Mean; SD = Standard Deviation; FAST = Functional Assessment Staging Test; MMSE = Mini Mental State Exam; CERAD= Consortium to Establish a Registry for Alzheimer's Disease neuropsychological battery. Compared to non-carriers, cognitively-unimpaired mutation carriers exhibited significantly worse CERAD word list delayed recall scores. Groups did not differ between in age, education, sex, FAST or MMSE scores. CERAD = Consortium to Establish a Registry for Alzheimer's Disease

### Segregated and integrated functional connectivity patterns differ between cognitively-unimpaired mutation carriers and non-carriers

We first examined whether there were distinct patterns of segregated and integrated functional connectivity between groups. Higher values in segregated functional connectivity maps denote that a given voxel has a greater number of strongly functionally connected links to other regions of the same functional network, whereas higher values in integrated functional connectivity maps mean that the voxel is an important hub for integrating information between brain communities.

Compared to non-carriers, mutation carriers exhibited less functional segregation and integration of posterior DMN regions that subserve episodic memory (Figure 1). Specifically, we observed less segregated and integrated functional connectivity in the precuneus, a hub of the DMN that is vulnerable to early tau accumulation in this cohort (24). We also found less integrated functional connectivity in the retrospenial cortex, which has been suggested to link subcortical systems (i.e., medial temporal regions) and cortical regions of the DMN (27). This suggests that the precuneus is functionally disconnected from regions within network as well as other brain communities, while the retrospenial cortex is disconnected from regions across networks.

**Figure 1.**
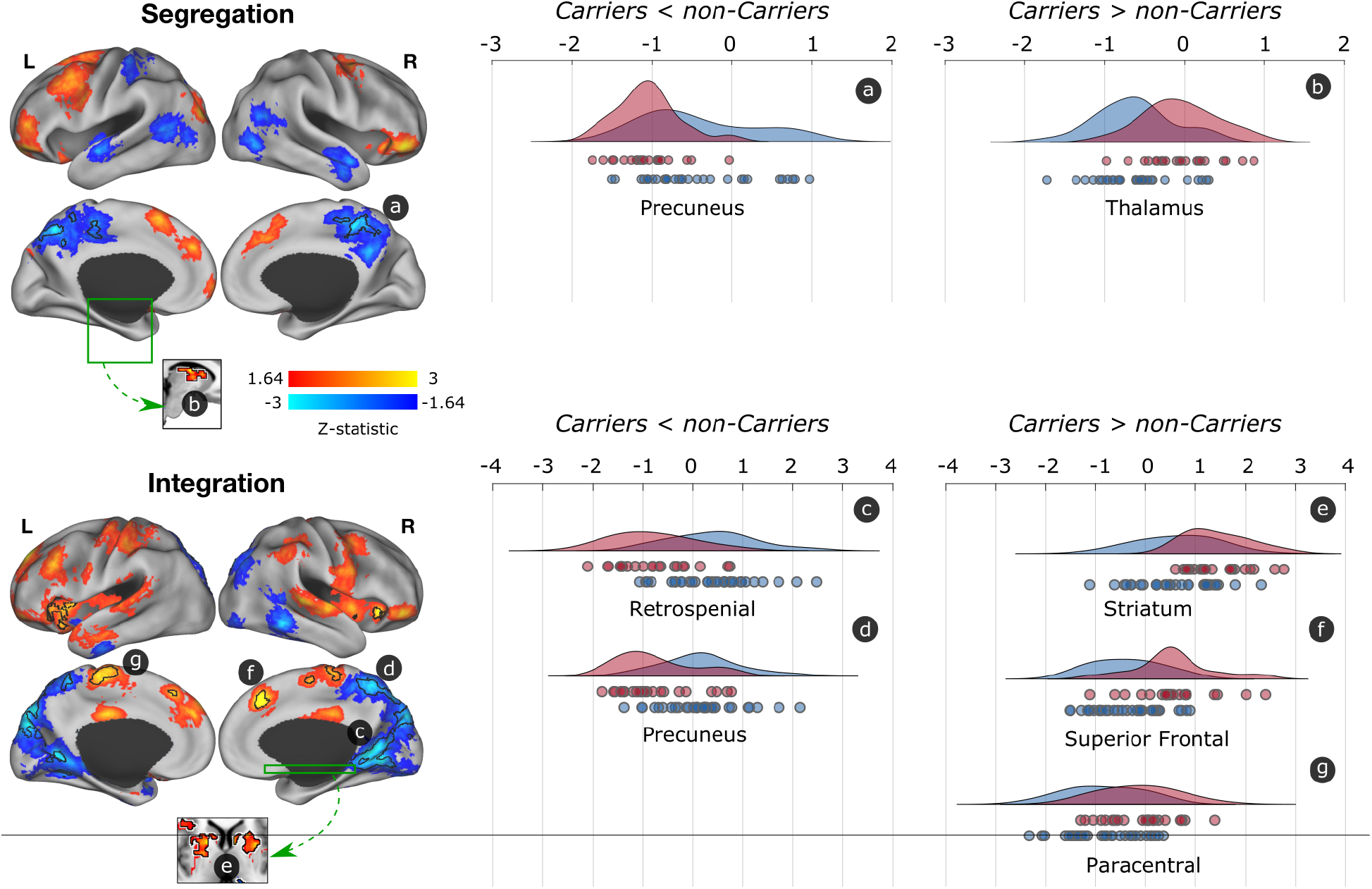
Differences in segregated and integrated network connectivity between cognitively-unimpaired mutation carriers and non-carriers. Differences in functional segregation and integration between mutation carriers and non-carries are displayed. Red-yellow colors show the trend (Z>1.64; p<0.1) of greater functional connectivity while blue regions display less functional connectivity in mutation carriers compared to non-carriers. Regions with a p<0.05 and surviving to multiple comparisons are indicated with black borders. The graphs at the right display the distribution of functional connectivity for the surviving clusters, with non-carriers being represented in blue and mutation carriers in red. Circles below the curves for each brain structure represent data for a single subject

We then conducted two seed-based post hoc analysis of the fMRI data to better characterize functional connectivity patterns of these two regions (see supplementary information). The precuneus seed exhibited less functional segregation with other DMN regions, including the posterior cingulate cortex, supramarginal, and temporal and medial frontal cortex. On the other hand, the retrosplenial cortex exhibited less functional integration with regions from the salience network, as well as the hippocampus and brainstem (Figure S1). Notably, the observed dysconnectivity of these structures with other brain regions is unlikely due to atrophy, a marker of neurodegeneration, as we did not observe significant atrophy in any of the posterior regions (Figure S2; see supplementary information). This suggests that the observed functional changes precede neurodegeneration and may instead be associated with other pathophysiological processes, such as accumulation of tau pathology, as we report in the next section.

Mutation carriers also exhibited greater segregated and integrated functional connectivity of mostly salience network regions compared to non-carriers (Figure 1). Specifically, we observed greater segregation in the thalamus, and integration in the striatum, medial frontal cortex, and paracentral lobule. This suggests that the thalamus is hyperconnected to within-network regions, and the other regions to other brain communities.

### Amyloid-β and tau burden are distinctively associated with functional connectivity maps in cognitively-unimpaired mutation carriers

We then examined the association between functional segregation and integration patterns with AD progression in the preclinical stage. Specifically, we conducted a novel bipartite graph analysis to elucidate the relationship between pathology burden in voxels that harbor high levels of pathology and the fMRI voxels with altered functional segregation or integration (Figure 2A).

**Figure 2.**
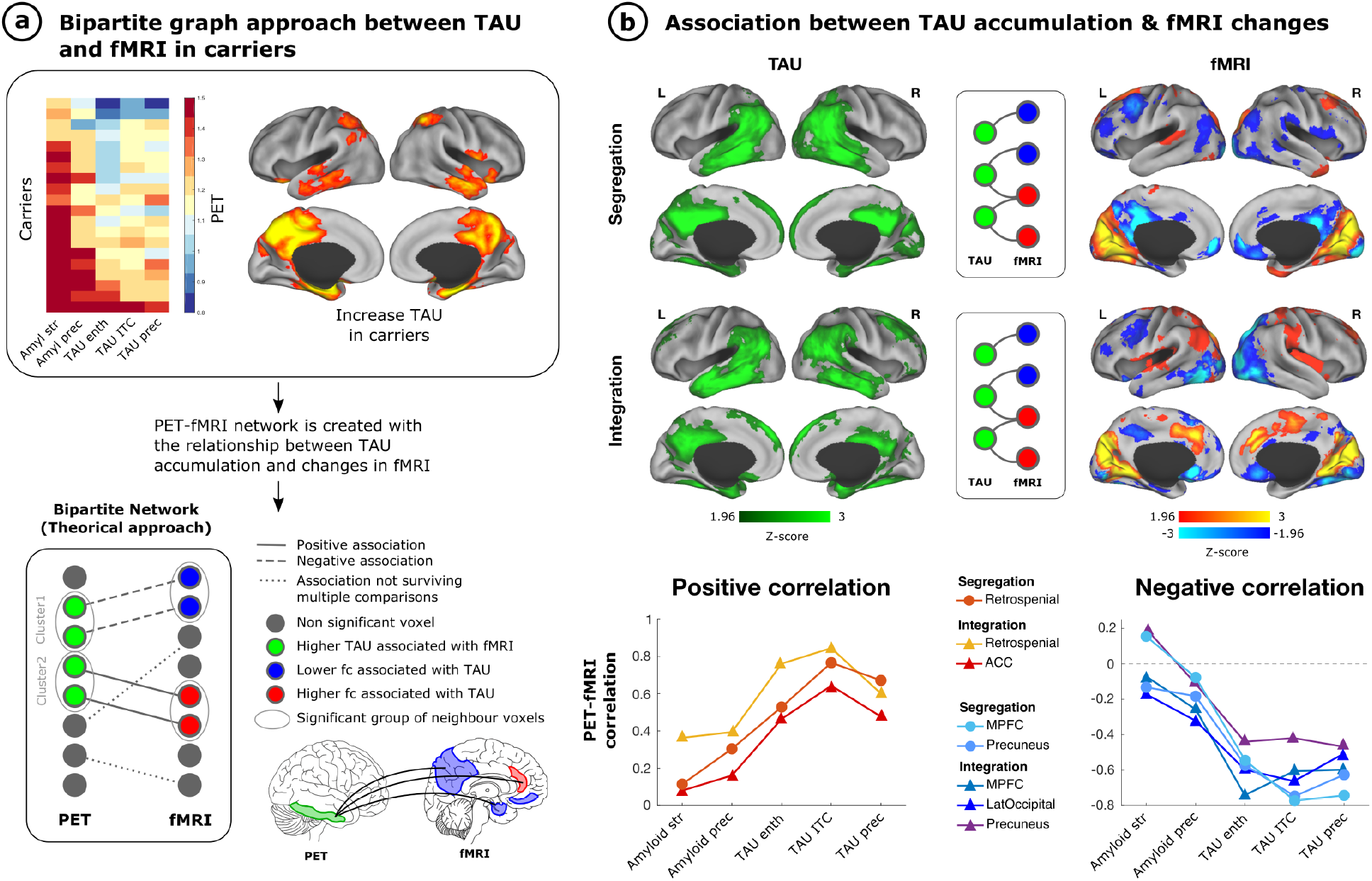
Relationship of segregated and integrated network connectivity, with amyloid-β and tau pathology in mutation carriers. We used a bipartite graph theory approach to investigate the association between whole-brain PET imaging and resting-state fMRI in mutation carriers. The top section of panel (a) illustrates the accumulation of amyloid-β and tau pathology in different brain regions that have been used to track disease progression in mutation carriers, together with a brain representation of the brain regions that exhibit tau accumulation the earliest in mutation carriers. The bottom part of panel (a) shows the bipartite graph approach that we employed on this study to test the relationship between tau burden and functional connectivity patterns. Specifically, we examined the association between voxel clusters with high tau burden and the fMRI voxels with altered functional segregation or integration. Panel (b) to the left exhibits brain regions with greater tau burden in mutation carriers compared to noncarriers, shown in green. Panel (b) shows fMRI voxels associated with greater tau burden that survived multiple comparisons for functional segregation and integration. Red-yellow colors represent greater functional connectivity while blue regions represent less functional connectivity. Graphs at the bottom of panel (b) illustrate the correlation of amyloid-β and tau pathology in regions that are most vulnerable to early pathology accumulation with brain regions that exhibited less or greater segregation or integration. ACC = anterior cingulate cortex; amyl = amyloid-β; enth = entorhinal cortex; fc = functional connectivity; ITC = inferior temporal cortex; LatOccipital = lateral occipital cortex; MPFC = medial prefrontal cortex; prec = precuneus; str = striatum

First, we found that the precuneus, inferior temporal and entorhinal cortices had the greatest accumulation of tau pathology in mutation carriers compared to non-carriers (Figure 2A), as previously reported(24). We then examined how tau burden in these regions related to segregated and integrated functional connectivity in mutation carriers. We found that greater tau burden in these brain structures was associated with less segregated and integrated functional connectivity of DMN regions, particularly the precuneus and medial prefrontal cortex, as well as the lateral occipital cortex (Figure 2B). That is, greater tau burden was related to a greater disruption and reduction in connectivity of these regions within network and across brain communities.

In turn, more tau accumulation was associated with greater segregated and integrated functional connectivity in the retrospenial cortex, and integrated functional connectivity in the anterior cingulate cortex (ACC), a hub of the salience network (Figure 2B). This suggests that, as tau accumulates, the retrospenial cortex becomes highly connected to regions within network and across brain communities, while the ACC becomes more functionally connected to regions within network only.

We then proceeded to examine the relationship between the functional segregation and integration of each of these regions (i.e., precuneus, medial prefrontal cortex, lateral occipital cortex, retrospenial cortex, and the ACC), with amyloid-β and tau pathology. Since amyloid-β pathology accumulates in the cortex more than a decade before tau does on this kindred, looking at the relationship between amyloid-β and tau with the functional segregation and integration of these regions may provide insight into how functional connections change with disease progression. Results suggest that there is greater functional segregation and integration of the retrospenial cortex, and integration of the ACC in the presence of tau pathology compared to amyloid-β burden (Figure 2B), while the opposite is true for the segregation and integration of DMN regions.

### Functional connectivity alterations are associated with worse episodic memory in cognitively-unimpaired mutation carriers

Finally, we investigated whether functional brain maps were related to the CERAD word list delayed recall scores in mutation carriers. We found that higher delayed recall scores were significantly associated with higher segregated functional connectivity in the left middle frontal gyrus, and lower functional connectivity in subcortical regions (Figure 3). We also observed that higher delayed recall scores were related to lower integrated functional connectivity in the precuneus and medial prefrontal cortex, and with greater integrated functional connectivity in the precentral cortex. We then examined whether functional connectivity mediated the relationship between pathology and delayed recall scores, or whether pathology mediated the relationship between functional connectivity and delayed recall scores. We did not find a relationship as none of the regions survived correction for multiple comparisons.

**Figure 3.**
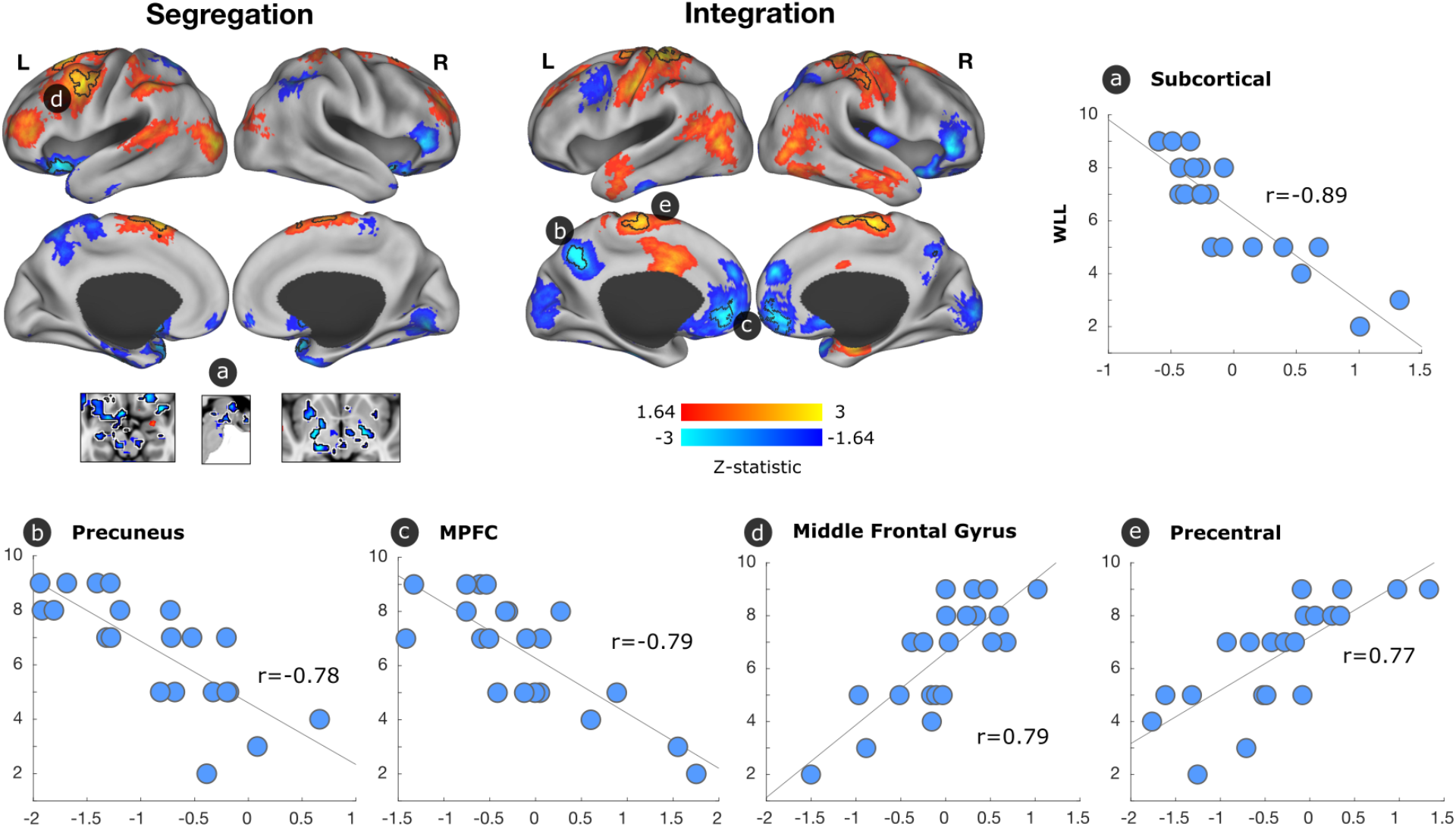
Association between segregated and integrated network connectivity, and word list delayed recall in mutation carriers. We conducted a general linear model to evaluate the functional connectivity patterns associated with word list delayed recall scores in mutation carriers. Regions that were associated but did not survive multiple comparisons (Z>1.64; p<0.1) are displayed on the brain, with black borders indicating regions that survived multiple comparisons (p<0.05). Red-yellow colors illustrate an association between better delayed recall scores and greater functional connectivity, and blue colors a relationship between better delayed recall scores and less functional connectivity. Scatterplots illustrate the relationship of word list delayed recall scores with functional connectivity in regions that survived multiple comparisons. MPFC = medial prefrontal cortex

## DISCUSSION

We provide novel *in vivo* evidence that amyloid-β and tau pathologies relate to distinct patterns of functional segregation and integration that resemble the patterns of pathology accumulation in AD and relate to episodic memory in preclinical ADAD. We found that posterior DMN regions, particularly the precuneus, one of the earliest sites of pathology accumulation in ADAD, and the retrospenial cortex, exhibited significantly less functional segregation and integration of networks compared to age-matched non-carriers. This decrease in functional segregation and integration of the precuneus with other regions within network and across brain communities was also strongly associated with greater tau burden in the entorhinal cortex, precuneus and inferior temporal gyrus, and less to amyloid-β, suggesting that dysconnectivity of the precuneus occurs closer to symptom-onset and may play a pivotal role in predicting disease progression early. Similarly, the medial prefrontal cortex, which is part of the DMN, exhibited less segregated and integrated functional connectivity in relation to greater tau burden in the same regions.

Both the precuneus and medial prefrontal cortex are core hubs of the DMN and support complex cognition and behaviors such as episodic memory(28–30). Prior studies have suggested that individuals at high risk for AD exhibit hyperconnectivity in the DMN, before a reduction in functional connectivity is observed(15). In fact, we previously reported that children ages 9 to 17 who carry the E280A mutation exhibited increased functional connectivity of DMN regions compared to age-matched non-carriers(31). This hyperconnectivity phase has been most strongly associated with amyloid-β pathology and the hypoconnectivity phase most strongly related to tau pathology burden(15). Prior studies with other ADAD mutations have also reported decreased functional DMN segregation as individuals approach symptom onset. One particular study demonstrated decreased functional connectivity of the precuneus and other posterior DMN regions in cognitively-unimpaired mutation carriers(32), while another showed that both anterior and posterior regions of the network are disrupted early(33). Notably, other studies have only observed less functional connectivity, including in the DMN and between networks, in cognitively impaired mutation carriers as measured by a score of 1 on the clinical dementia rating (CDR) scale, and not in cognitively unimpaired mutation carriers (CDR=0)(11, 13). These studies measured functional connectivity by either calculating a global resting-state functional connectivity signature(11) or composite scores comprised of regions chosen a priori(13). These findings suggest that the voxel-wise connectivity analyses conducted in this study provide high-resolution connectome information that increases the sensitivity to detect subtle disease-related changes that occur in the preclinical stage of AD.

Similar to the DMN, the salience network has also been shown to undergo a phase of hyperconnectivity associated with mostly amyloid-β, followed by hypoconnectivity(15). Consistent with these findings, mutation carriers exhibited greater segregated functional connectivity in regions connected to the salience network such as the thalamus, and greater functional integration in the striatum. Similarly, greater tau burden was related to greater functional integration of the ACC, a salience network hub. While some studies have not observed changes in the salience network in preclinical AD, others have suggested that the salience network becomes disrupted early in ADAD(33) and that it is particularly vulnerable to early AD-related changes together with the DMN. Our findings suggest that some salience network regions (ACC and striatum) are hyperconnected to other brain communities in the preclinical stage of AD, and others (thalamus) to within network. It is possible that the observed segregated hyperconnectivity serves as a predictor for future pathology accumulation as animal studies have demonstrated that neuron-to-neuron propagation of tau pathology is facilitated by greater synaptic connectivity and neuronal activity(21, 34). The same can be said for the retrospenial cortex, which is strongly connected to medial temporal regions that subserve episodic memory. Perhaps these connections will weaken with time, when tau aggregates in these regions as most of them are primarily affected by amyloid-β at this stage.

In regards to the relationship between functional connectivity and episodic memory, we observed that less integrated functional connectivity in the precuneus and medial prefrontal cortex was associated with better memory test scores. That is, mutation carriers with better episodic memory exhibited hypoconnectivity of these two DMN regions when integrating information with other networks of the brain. We also found a significant relationship between greater functional integrity in the left middle frontal gyrus and better episodic memory. While these findings may appear to conflict with the observed patterns in functional connectivity associated with pathology at a first glance, prior research suggests that hyperconnectivity in the left frontal cortex may represent a compensatory mechanism or form of neural reserve. Specifically, in a study examining individuals with ADAD and sporadic AD researchers reported that greater functional connectivity in the left frontal cortex was associated with less cognitive decline and global cognition, as well as a diminished effect of tau on cognitive performance in the presence of greater left frontal cortex connectivity in the early stages of AD(35). As such, we hypothesize that the observed hyperconnectivity of the left frontal cortex associated with better episodic memory may reflect a form of compensation or neural reserve in the preclinical stage of AD.

Altogether, our findings suggest that in the presence of tau pathology, mutation carriers exhibit largely dysconnectivity of DMN brain regions, particularly posterior regions, and increased connectivity of structures that are associated with the salience network and others that are critical for integrating information across neural systems. Based on what it has been reported and our own findings, one could hypothesize that elevated amyloid-β levels contribute to an increase in functional connectivity or neuronal firing between the precuneus and other DMN regions, and later for salience networks regions, increasing the vulnerability of these regions for early tau deposition in ADAD, and later becoming disconnected in the presence of tau pathology, further contributing to cognitive decline. Studying the relationship between AD-related pathology and the functional organization of the brain in younger mutation carriers, as well as longitudinally, may provide the opportunity to test the aforementioned hypothesis.

The current study has multiple strengths. First, we did not rely on presenting symptoms or predicted risk (e.g., based on amyloid-β PET levels or APOE genotype) to infer whether individuals will develop dementia. Instead, we tested our hypotheses in individuals who have a well-characterized disease and clinical trajectory. In addition, we examined *in vivo* amyloid-β and tau pathology using PET imaging, which is considered the gold standard for quantifying and mapping brain pathology in AD. Mutation carriers were also young and otherwise generally healthy, which minimizes potential age-related confounding variables that contribute to brain dysfunction and cognitive decline (e.g., cardiovascular disease). Further, we conducted voxel-level connectivity analyses, which provide high-resolution connectome information that increases the sensitivity to detect subtle disease-related changes that occur in the preclinical stage of AD. Finally, the very homogeneous clinical profile of mutation carriers allows us to infer how functional connections may change as the disease progresses.

The present study also has caveats that must be considered when interpreting the data. First, our sample size is relatively small compared to other studies of AD. However, individuals with these mutations are relatively rare and all our participants carry a single mutation, which makes our sample highly homogeneous compared to other cohorts, and one of the largest single mutation ADAD samples with PET imaging and fMRI. Nonetheless, findings should be validated on larger ADAD samples in the future. Further, more research is needed to examine whether our findings in ADAD generalize to preclinical late-onset AD, as well as whether other genetic risk factors known to be important in may impact the results in this kindred. We are currently conducting the first longitudinal biomarker study with this cohort, which will provide greater insight into how annual change in functional connectivity relates to *in vivo* pathology burden and cognitive decline over time.

Altogether, these findings highlight the importance of not limiting analyses to examining group averages or specific seeds, as it can obscure the unique and often subtle patterns of functional disintegration associated with disease progression. Our data also underscores that more attention must be given to the precuneus as an early marker of AD and key player in functional dysconnectivity. Findings enlighten our understanding of how AD-related pathology may distinctly alter the functional architecture of the brain to interfere with the integration of information within and across different neural systems and possibly propagate the spread of pathology, ultimately resulting in cognitive impairment and dementia.

## MATERIALS AND METHODS

### Study design and participants

*PSEN1* E280A carriers and age- and education-matched noncarriers from the Massachusetts General Hospital COLBOS (Colombia-Boston) longitudinal biomarker study participated in this study. Exclusion criteria included a history of psychiatric disorders, illiteracy, stroke, epilepsy, traumatic brain injury, kidney failure, human immunodeficiency syndrome, or substance abuse. To be included in this study participants had to demonstrate no cognitive impairment on a standard cognitive battery, including a clinical diagnostic rating scale (CDR) score of 0, a Functional Assessment Staging Test (FAST) score of 2 or less, and a MMSE score of 26 or greater. Demographic information is presented in Table 1.

The study was approved by both the institutional ethics review boards of the University of Antioquia in Colombia and Massachusetts General Hospital in Boston. Participants provided signed informed consent before participating in any procedures.

All participants in this study travelled from Colombia to Boston (USA) and underwent PET imaging and MRI at the Massachusetts General Hospital. Participants and investigators were blind to the genetic status of the individuals.

### Genotyping

Genomic DNA was extracted from blood using standard protocols. *PSEN1* E280A characterization was done at the University of Antioquia as previously described(36). Genomic DNA was amplified with the primers PSEN1-S 5’ AACAGCTCAGGAGAGGAATG 3’ and PSEN1-AS 5’ GATGAGACAAGTNCCNTGAA 3’. We used the restriction enzyme BsmI for restriction fragment length polymorphism analysis. Each participant was classified as a *PSEN1* E280A carrier or non-carrier.

### Neuropsychological measures

A Spanish version of the Consortium to Establish a Registry for Alzheimer’s Disease (CERAD) word list learning test that is sensitive to early episodic memory changes in mutation carriers was used in this study (37). On this test participants were required to learn 10 items over 3 trials. Participants were then asked to recall as many words as they could from the previously learned list after a 10-minute delay (delayed recall score). Higher scores on the delayed recall represent better performance on the memory task.

### MRI and PET data acquisition and preprocessing

See Supplementary Information for acquisition and preprocessing procedures for MRI, and 11C Pittsburg compound B and [F18] Flortaucipir PET.

### Integration and segregation analysis

The human brain exhibits a modular organization where different subsets of voxels form communities with specialized tasks(38). Based on this modular organization, the brain’s functional connectivity can be separated in: i) segregated links connecting all the voxels belonging to the same community; and ii) integrated links connecting links of each voxel with the rest of the communities of the brain for higher cognitive integration.

A whole brain functional connectivity matrix was computed using Pearson *r* correlation coefficient in the time series of each pair of voxels. A 49,314 x 49,314 association matrix was obtained for each participant and negative values were removed(14, 16). The 49,314 grey matter voxels included cortical, subcortical, brainstem and cerebellum regions. We generated a whole brain seven resting-state network parcellation to classify each link in the connectivity matrix as segregation or integration. The seven resting-state networks of the Yeo parcellation were used as initial seed to extend these networks to also cover subcortical, brainstem and cerebellar regions(39).

We generated a parcellation scheme taking advance of the same acquisition scanner and sequences offered by our local database Brain Genomics Superstruct Project Open Access (GSP)(40) based on a sample of 100 participants. Individual-subject seed connectivity was computed for visual, sensory-motor, dorsal attention (lateral-visual), ventral attention (salience), limbic, frontoparietal and default mode networks. The average population map of each network was calculated, and each voxel of the atlas was assigned to each network based on the maximum probability of each voxel belonging to each of the networks (Figure S3). If the link’s start and end voxels belonged to the same resting state network, the link was then classified as a segregated link; otherwise, the link was classified as an integrated link. For each participant, we computed the weighted degree of segregated and integrated links separately obtaining two brain maps, one for each connectivity type(41). The weighted degree represents the strength of connections that each voxel in the brain has. Higher values in the segregated map means that a given voxel has a high number of strong connecting links to other regions of the same functional network. On the other hand, high values in the integrated map means that this voxel is an important hub for integration of information between brain communities.

### Statistical Analyses

#### Demographics and cognition

We conducted independent samples t-tests using SPSS Statistics, version 24.0 (Armonk, NY: IBM Corp.) to examine group differences in demographic and cognitive variables. We used Chi-squared test to examine sex and FAST differences between groups. Analyses used a family-wise significance threshold of *p* < 0.05 to correct for multiple comparisons and Cohen’s *d* to calculate effect sizes.

#### Segregation and integration group differences

We used a general linear model to compute a two-sample t-test to compare segregation and integration weighted degree maps between the mutation carriers and the non-carriers, covarying for motion (MATLAB V9.6, The MathWorks, Inc., Natick, MA, USA). Whole-brain correction for multiple comparisons was applied using Monte Carlo simulation with 10,000 iterations to estimate the probability of false positive clusters with a two-tailed p < 0.05 (3dClustSim, afni.nimh.nih.gov).

#### Association between resting-state fMRI and PET images

We used a novel bipartite graph approach to examine the association between amyloid-β and tau burden and functional segregation and integration in mutation carriers (Figure 2). Since the bipartite graph approach is computationally complex, we downsampled PET and fMRI segregation and integration maps to 6mm (6,779 voxels). We employed a linear general model, adjusting for motion in fMRI, to measure the relationship between PET burden in a specific voxel, and the segregation and integration in another voxel, for each pair of PET and fMRI voxels. We obtained four different association matrices: i) relationship of tau with functional segregation; ii) relationship of tau with functional integration; iii) relationship of amyloid-β with functional segregation; and iv) relationship of amyloid-β with functional integration. We then adapted the cluster-level Monte Carlo multiple comparison correction technique to bipartite networks to control for type I errors of bipartite graphs at the link level. Only clusters of PET voxels (group of contiguous voxels with significant association) that connected to cluster of fMRI voxels with a minimum size survived multiple comparison. To estimate the minimum size of a PET cluster and fMRI cluster to be significant with a p < 0.05 we generated 10000 bipartite random networks with the same smoothing properties. We computed the likelihood of significant cluster sizes due to chance in the generated random network and only those with a p < 0.05 were reported. The endpoints of the links that survived multiple comparisons are displayed in both PET and fMRI modality in Figure 2.

#### Functional integration and segregation association with episodic memory

We conducted a general linear model to examine the relationship between weighted degree maps in mutation carriers and the CERAD word list delayed recall, covarying for motion (MATLAB V9.6, The MathWorks, Inc., Natick, MA, USA). Whole-brain correction for multiple comparisons was applied using Monte Carlo simulation with 10,000 iterations to estimate the probability of false positive clusters with a two-tailed p < 0.05 (3dClustSim, afni.nimh.nih.gov).

## ACKNOWLEDGMENTS

The authors thank the Colombian families for contributing their valuable time and effort, without which this study would not have been possible. We also thank Francisco Piedrahita, Alex Navarro, Yamile Bocanegra and Claudia Ramos from Grupo de Neurociencias, Universidad de Antioquia in Medellín, Colombia, as well as Heirangi Torrico-Teave, Arabiye Artola, Jairo Martínez, and Diana Munera from the Massachusetts General Hospital in Boston, MA, for helping coordinate visits to Boston and assisting with data collection and processing.

Dr. Guzmán-Vélez was supported by the National Institutes of Health (NIH) National Institute on Aging (NIA) (K23AG061276). Dr. Quiroz was supported by the NIH NIA (R01 AG054671). Mr. Fox-Fuller reports NRSA support from the NIH NIA (F31AG06215801A1). Dr. Vila-Castelar was supported by a grant from the Alzheimer’s Association (2019A005859). Dr. Pardilla-Delgado was supported by the NHLBI (3T32HL007901-19S1). Dr. Sepulcre was supported by grants from the NIH (R01 AG061811, R01 AG061445). Dr. Schoemaker received postdoctoral fellowships from the American Heart Association (20POST35110047) and the Fonds de recherche du Québec - Santé (254389).

