## Supplemental Information for "Amyloid-β and tau pathologies relate to distinctive brain dysconnectomics in preclinical autosomal dominant Alzheimer’s disease"

### SUPPLEMENTARY INFORMATION

#### Methods

**MRI data acquisition and preprocessing.** The MRI image acquisition was performed on a Siemens 3T Tim Trio system using a 12-channel phased-array head coil. A high-resolution 3D T1-weighted magnetization prepared rapid gradient-echo (MPRAGE) sequence was used with the following parameters: 1mm isotropic voxels; 160 sagittal slices; acquisition matrix size=256×256; repetition time (TR)=2300 ms; echo time (TE)=2.98 ms; field of view (FOV)=256 mm. To measure blood oxygenation level dependent (BOLD) signal a functional gradient-echo echo-planar pulse sequence was used with the following parameters: TR of 3 seconds, flip angle 858, TE 30 ms, matrix 72 3 72, field of view 216 x 216 mm, 47 x 3 mm axial slices, which resulted in isotropic voxels of 3mm and 124 volumes lasting 6 minutes. Participants were asked to lie flat, keep eyes open, and remain awake. Bi-temporal foam pads were used to restrict head motion.

MRI data was preprocessed using FMRIB Software Library v5.0.7 (FSL) and MATLAB R2019b. The anatomical T1 preprocessing pipeline included: reorientation to right-posterior-inferior (RPI); alignment to anterior and posterior commissures; skull stripping; gray matter, white matter and cerebrospinal fluid segmentation; as well as a computation of non-linear transformation between individual skull-stripped T1 and 2mm resolution MNI152 template images. The functional MRI preprocessing pipeline included: slice timing correction; reorientation to RPI; rigid-body realignment of functional volumes within runs (6 parameters linear transformation); computation of the transformation between individual skull-stripped T1 and mean functional images using linear boundary based registration (https://fsl.fmrib.ox.ac.uk/fsl/fslwiki/FLIRT_BBR); intensity normalization; removal of confounding factors from the data using linear regression that included 12 motion-related covariates (rigid motion parameters and its derivatives), linear and quadratic terms, and five components each from the lateral ventricles and white matter. Global signal regression was not applied due to the spurious correlations this can introduce. Non-linear transformation of resting-state data to MNI space, concatenating the transformation from functional to structural and from structural to 3mm MNI standard space, spatial smoothing with an isotropic Gaussian kernel of 8mm FWHM, and band-pass filtering (0.01–0.08 Hz) to reduce low-frequency drift and high-frequency noise were also performed. Head motion was quantified using realignment parameters obtained during image preprocessing, which included 3 translation and 3 rotation estimates. Scrubbing of time points with excess head motion, Jenkinson frame displacement > 0.2mm, were interpolated. Most of the participants had 2 resting state fMRI acquisitions. We concatenated both preprocessed sequences and used the 120 time points with a frame displacement of <0.2mm (6 minutes). There was overall very little motion since all subjects were young and cognitively unimpaired. All participants had at least 120 time points and no subject needed to be discarded.  Nonetheless, we took a step further to ensure that the results were not contaminated by motion and added the mean motion of each subject as a cofounding factor in all statistical analyses. The distribution of the correlations across all-time series were inspected for possible noise contamination. We did not observe outliers from the whole-brain connectivity distributions across participants.

**11C Pittsburg compound B and [F18] Flortaucipir PET acquisition procedures.** Each individual PET data set was rigidly co-registered to the subject's MPRAGE MR data using SPM8 (Wellcome Department of Cognitive Neurology, Function Imaging Laboratory, London). As reported previously(1), [F18] Flortaucipir (FTP) images were acquired between 80 and 100 minutes after a 9.0 to 11.0 mCi bolus injection in 4 separate 5-minute frames. [F18] FTP specific binding was expressed in FreeSurfer (FS) ROIs as the standardized uptake value ratio (SUVR) to the cerebellum, as previously reported(2). The spatially transformed SUVR PET data was smoothed with a 8mm Gaussian kernel to account for individual anatomical differences(3). PET data were down-sampled to 6mm in order to compute the bipartite association between tau and fMRI.

11C Pittsburg compound B (PiB) PET was acquired with a 8.5 to 15 mCi bolus injection followed immediately by a 60-minute dynamic acquisition in 69 frames (12x15 seconds, 57x60 seconds). 11C PiB PET data were expressed as the distribution volume ratio (DVR) with cerebellar grey as reference tissue. Regional time-activity curves were used to compute regional DVRs for each region if interest (ROI) using the Logan graphical method applied to data obtained between 40 and 60 minutes after injection(4). Of note, we did not have 11C PiB PET data for one participant.

**Brain volume analysis.** An optimized voxel-based morphometry was used(5) with FSL(6) to examine volumetric differences between mutation carriers and non-carriers. The partial gray matter volume estimations were transformed to 2mm MNI 152 standard space using non-linear registration and the resulting images were averaged and flipped along the x-axis to create a left–right symmetric, study-specific gray matter template. All native gray matter images were then non-linearly registered to this study-specific template and modulated to correct for local expansion (or contraction) due to the non-linear component of the spatial transformation. Finally, the modulated gray matter images were smoothed with an isotropic Gaussian kernel with a sigma of 3mm.

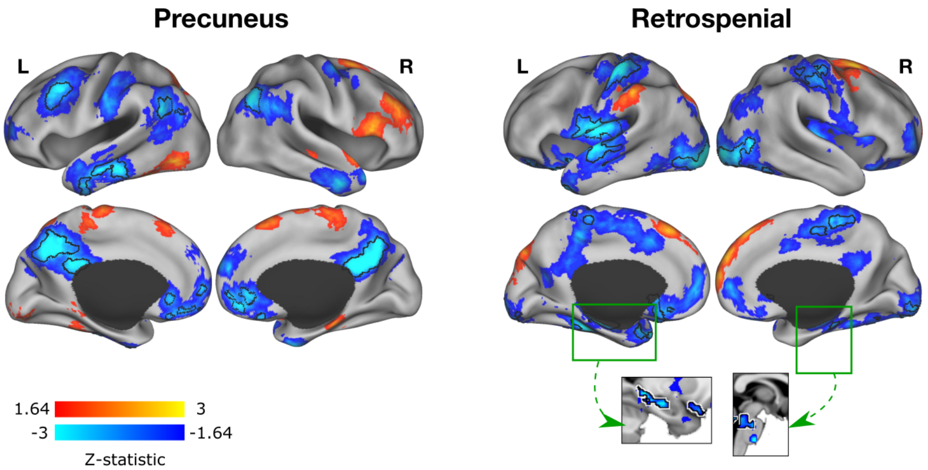


**Figure S1. Seed-based functional connectivity analyses with the precuneus and retrosplenial cortex.** We performed a two seed-based post hoc analysis of the fMRI data. We specifically seeded the precuneus, which showed significantly less functional segregation, and the retrospenial cortex, which showed significantly less functional integration, in mutation carriers compared to non-carriers. The precuneus seed exhibited less functional segregation to other DMN regions, including the posterior cingulate cortex, supramarginal, temporal and medial frontal cortex. In turn, the retrospenial cortex exhibited less functional integration with regions from the salience network, as well as the hippocampus and brainstem, all of which survived multiple comparison correction.

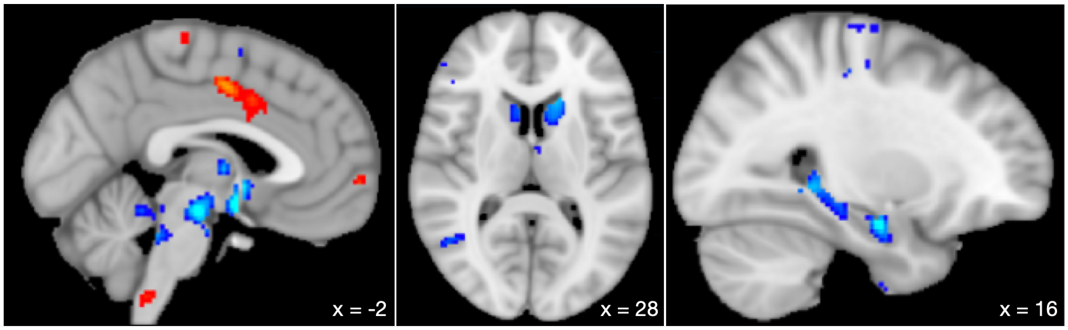


**Figure S2. Atrophy in mutation carriers vs non-carriers.** We used FSL voxel-based morphometry adjusting for head size to compare the grey matter volume of mutation carriers and non-carriers and provide a better understanding of whether atrophy in the precuneus could be leading to less functional connectivity of the precuneus. We did not find significant volume loss in the precuneus at this stage of the disease. However, we found greater volume loss in the hippocampus, caudate nucleus, and midbrain. Blue colors represent lower volume in mutation carriers and red/orange colors higher volume in mutation carriers.

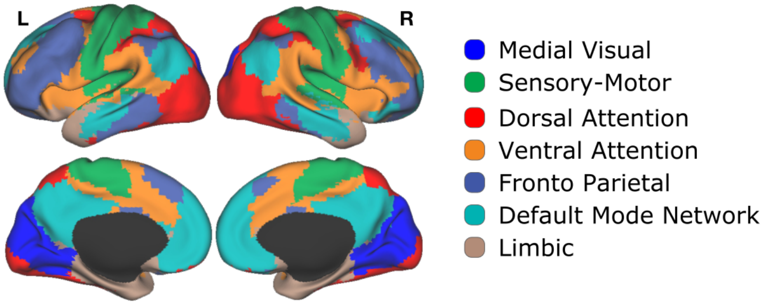


**Figure S3. Resting-state networks parcellation.** We generated this parcellation scheme taking advance of the same acquisition scanner and sequences offered by our local database Brain Genomics Superstruct Project Open Access (GSP). The parcellation scheme includes subcortical structures, as well as the brainstem and cerebellum.
